# Feasibility of using a fully immersive virtual reality system for kinematic data collection

**DOI:** 10.1101/547992

**Authors:** Kate A. Spitzley, Andrew R. Karduna

**Author notes:** Corresponding Author: Kate A. Spitzley.

## Abstract

Commercially-available Virtual Reality (VR) systems have the potential to be effective tools for simultaneous visual manipulation and kinematic data collection. Previously, these systems have been integrated with research-grade motion capture systems to provide both functionalities; however, they are yet to be used as stand-alone systems for kinematic data collection. This study aimed to validate the HTC VIVE VR system for kinematic data collection by evaluating the accuracy of its position and orientation signals. The VIVE controller and tracker were each compared to a Polhemus Liberty magnetic tracking system sensor for angular and translational measurement error and signal drift. A sensor from each system was mounted to opposite ends of a rigid segment which was driven through fifty rotations and fifty translations. Mean angular errors for both the VIVE tracker and controller were below 0.4°. Mean translational error for both sensors was below 3 mm. Drift in the Liberty signal components was consistently lower than drift in VIVE components. However, all mean rotational drift measures were below 0.1° and all mean translational measures were below 0.35 mm. These data indicate that the HTC VIVE system may be a valid and reliable means of kinematic data collection. However, further investigation is necessary to determine the VIVE’s suitability for capturing extremely minute or high-volume movements.

## 1. Introduction

Recent advances in Virtual Reality (VR) technology have expanded our ability to integrate immersive 3D visual environments with optical motion capture. Platforms such as Vicon Reality and OptiTrack for VR have combined their marker-based systems with VR headsets and controllers to augment both the VR experience and research capabilities (Vicon.com, Optitrack.com, 2018). However, these systems still require the use of research-grade motion capture cameras to collect kinematic data. Although research-grade systems provide robust measures of position and orientation, there are limitations due to their high price and low portability. While these may not be barriers for more established institutions, those working in underfunded programs, teaching institutions, classroom settings, and clinics may find them restrictive. The use of a VR system for simultaneous immersion in 3D virtual environments and kinematic data collection could provide the benefits of these integrated systems, the low price point of VR systems, and high portability that existing systems cannot offer.

The HTC VIVE Virtual Reality System (VIVE) and Oculus Rift systems are both prime candidates for this application as they are each available for under $1000 and extremely portable. However, the tracked area for the VIVE is about 5 m x 5 m while the tracked area for the Rift is about 1.5 m x 1.5 m, making the VIVE a more suitable option for kinematic data collection (Martindale, 2018). The VIVE system consists of two small light-emitting boxes (9 cm x 9 cm x 6 cm), two lightweight handheld controllers (0.31 kg), a round tracker (0.36 kg), and a lightweight fully immersive headset (0.47 kg). The shipping weight for this entire system is under 2 kg. The position and orientation of the tracker, headset, and controllers are tracked in real time, allowing for realistic motion feedback in to the virtual visual environment.

The goal of this study was to determine the accuracy of the VIVE handheld controller and tracker, two different configurations of VIVE sensors, in comparison to an industry gold standard motion capture system, the Polhemus Liberty (Liberty) magnetic tracking system. This system has been used extensively in basic and clinical research settings (Amasay and Karduna, 2013; Kahol et al., 2009, 2008; Kwon et al., 2012). The manufacturer reported static accuracy of this system is 0.15° and 0.76 mm, and independent validation studies have corroborated these claims, showing root mean square (RMS) values as low as 0.2 mm (Nafis et al., 2006; Polhemus, 2012). This system was chosen for comparison due to its’ high reported accuracy and frequent use for collecting kinematic data (Amasay and Karduna, 2013; Dadarlat et al., 2015; Kahol et al., 2009; Lin and Karduna, 2016; Nafis et al., 2006; Polhemus, 2012). Measures of static drift in position and orientation, static translation, and static rotation were compared between the VIVE controller and the Liberty sensor and the VIVE tracker and the Liberty sensor. Feasibility of using the VIVE sensors for kinematic data collection was determined by their accuracy when compared to the Liberty sensor.

## 2. Methods

### System Setup

The VIVE VR system was set up as detailed by the user’s manual in order to establish proper tracking of the sensors. The lighthouses were set 6 m apart, mounted directly to the laboratory wall at a height of 2.1 m, angled downward at an angle of 30°, and connected by a synchronization cable (“htc VIVE User Guide,” 2016). The VIVE headset was located on a stable, flat platform within the tracked space. The system’s room setup protocol was run before each data collection session. This protocol establishes the location of the floor and the play area (the area in which the sensors will be tracked). The Liberty magnetic tracking system (Polheums, Colchester, VT) was set up according to the user’s manual and best practices (Polhemus, 2012). The Liberty transmitter was placed approximately 0.5 m from the sensor and metal was cleared from the space to eliminate interference (Figure 1).

**Figure 1.**
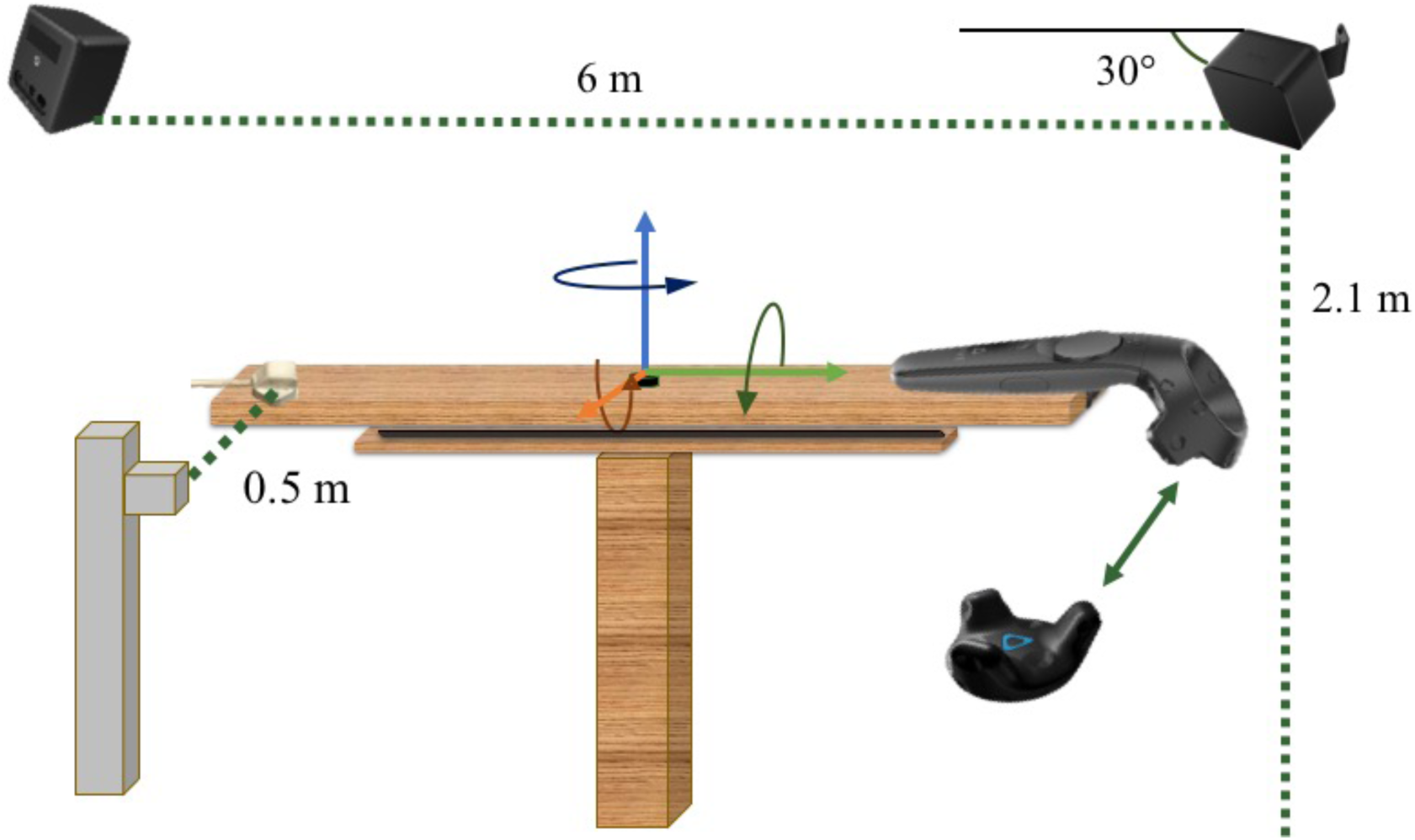
HTC VIVE lighthouse boxes mounted 2.1 m high, 6 m apart, tilted downward by 30 °, as suggested in the user guide. Polhemus Liberty base station positioned 0.5 m from the tracker. VIVE and Liberty sensors mounted to opposite ends of a rigid segment with six degrees of freedom. Graphics not to scale.

### Experimental Protocol

The VIVE sensor (tracker or controller) was mounted to a rigid segment opposite a sensor from the Liberty magnetic tracking system; the segment was then mounted to a ball-and-socket fixture, which was set on a linear gear track. This setup allowed for 360° of rotation about three axes and 61 cm of translation along three axes. The segment holding both sensors was moved through fifty rotations about each axis and fifty translations along each axis of the respective sensor (Figure 2). In total, 300 ten-second samples were collected while the segment was held static at each increment of motion. Increments ranged from 0 – 50° and 0 – 30 cm. All tests were completed first with the VIVE controller and Liberty sensor, then with the VIVE tracker and Liberty sensor. Testing order was chosen at random. The rig holding the sensors was stationed in the center of the VIVE system’s tracked space. Both systems sampled at a rate of 120 Hz. Data were collected simultaneously from the sensors using a custom Unity program (Unity Technologies, San Francisco, CA, USA) and analyzed using a custom LabVIEW program (National Instruments, Austin, TX, USA).

**Figure 2.**
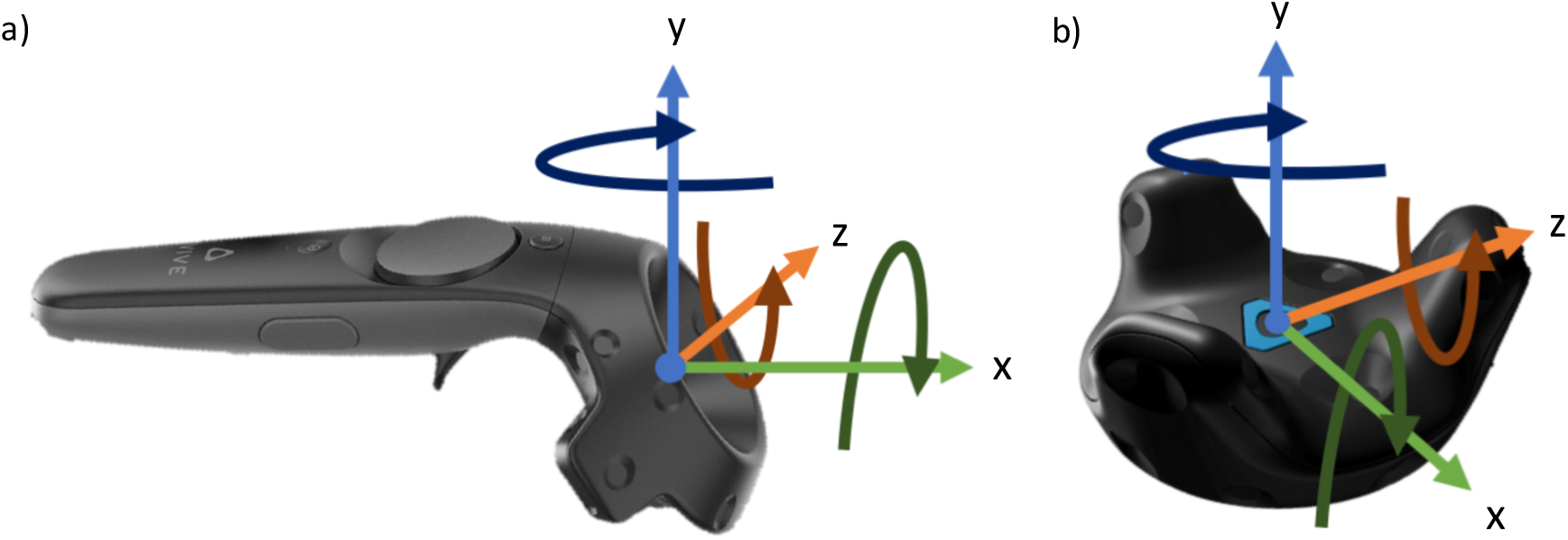
Coordinate systems of the HTC VIVE controller (a) and tracker (b) overlaid on their respective models.

### Data Reduction and Analysis

Translational error between the two systems was determined by taking the mean position over each ten-second period and using the distance formula to quantify the translation of each sensor from sample to sample. The distance measured by the VIVE was then subtracted from the distance measured by the Liberty. As neither system reported consistently higher or lower values than the other, indicating random error, the absolute value of the difference in measurement was used in order to avoid erroneously low error values.

Rotational error between the two systems was determined by comparing the change in helical angle measurement in each sensor from sample to sample. Helical angles were chosen because the global coordinate systems of the VIVE and Liberty systems were not aligned. The mean of each orientation component was taken over the ten-second collection period. The averaged Liberty components were decomposed from their Euler sequence using the system’s reported attitude matrix (Polhemus, 2012). Helical angle change from sample to sample was calculated using these mean raw rotational components from each system (Spoor and Veldpaus, 1980). The helical angle change was compared between the two systems by subtracting the change measured by the VIVE from the change measured by the Liberty. Again, random error was seen and as a result, the absolute value of these differences was used.

Drift in all six signal components was quantified using RMS of each ten second static collection period. The mean value of each signal component was first subtracted from each sample in that signal component; the remainders were used to determine RMS values for each signal component.

## 3. Results

The VIVE tracker and controller showed a mean rotational error of 0.13 ± 0.08° and 0.3 ±0.07°, respectively (Figure 3a, b). Mean translational error for the tracker and controller was 1.7 ± 0.4 mm and 2.0 ± 0.8 mm, respectively (Figure 3c, d). When tested together, the tracker and Liberty sensor showed mean rotational drift of 0.06 ± 0.07° and 0.003 ± 0.000°, respectively (Figure 4a) and mean translational drift of 0.27 ± 0.13 mm and 0.02 ± 0.00 mm, respectively (Figure 4c). When tested alongside the controller, the controller and Liberty sensor showed mean rotational drift of 0.01 ± 0.00 ° and 0.0002 ± 0.0001°, respectively (Figure 4b) and mean translational drift of 0.28 ± 0.13 mm and 0.01 ± 0.00 mm, respectively (Figure 4d).

**Figure 3.**
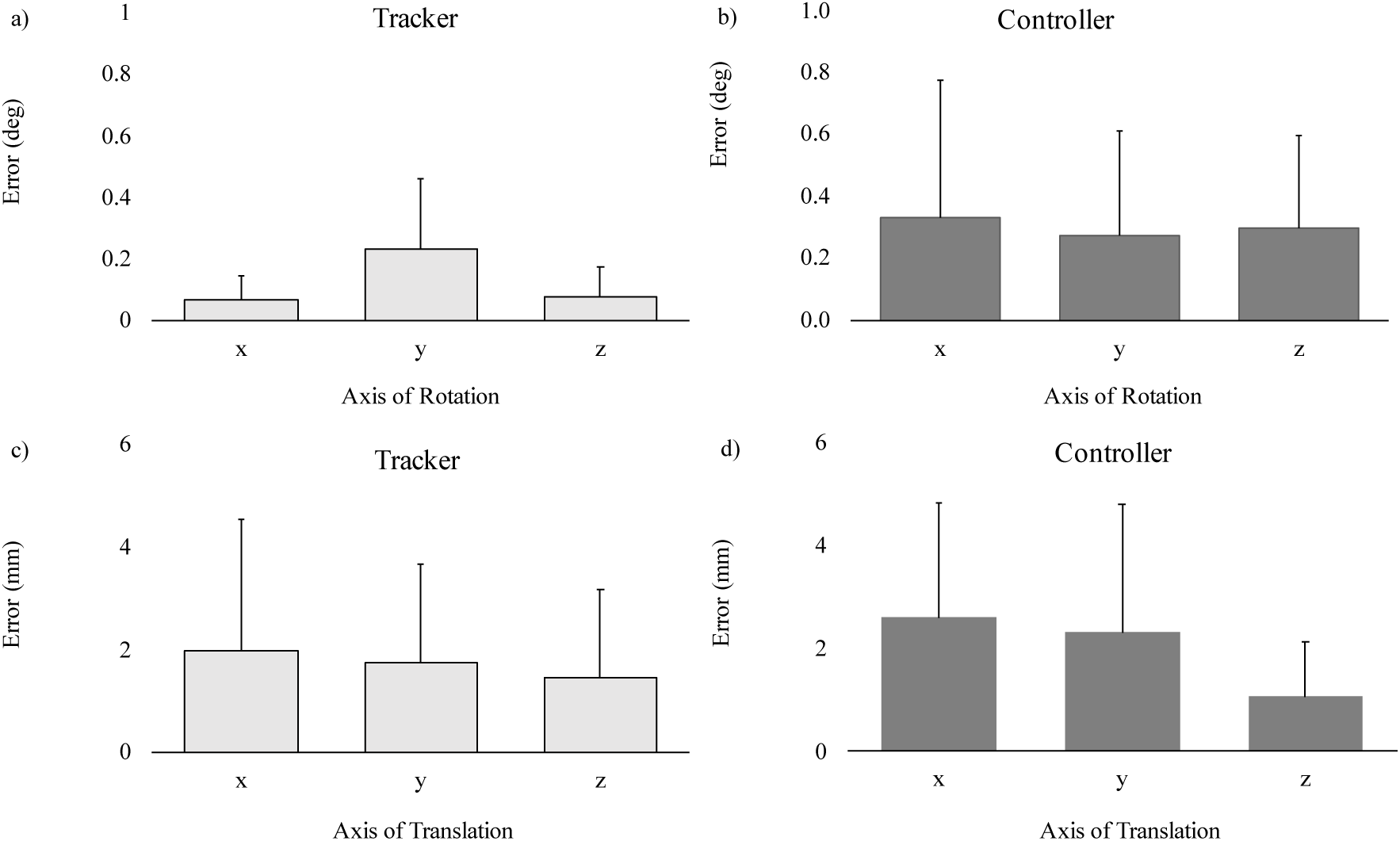
Comparisons of rotational (a and b) and translational (c and d) measurement errors (mean ± SD) between the VIVE tracker and Liberty sensor (a and c) and the VIVE controller and Liberty sensor (b and d).

**Figure 4.**
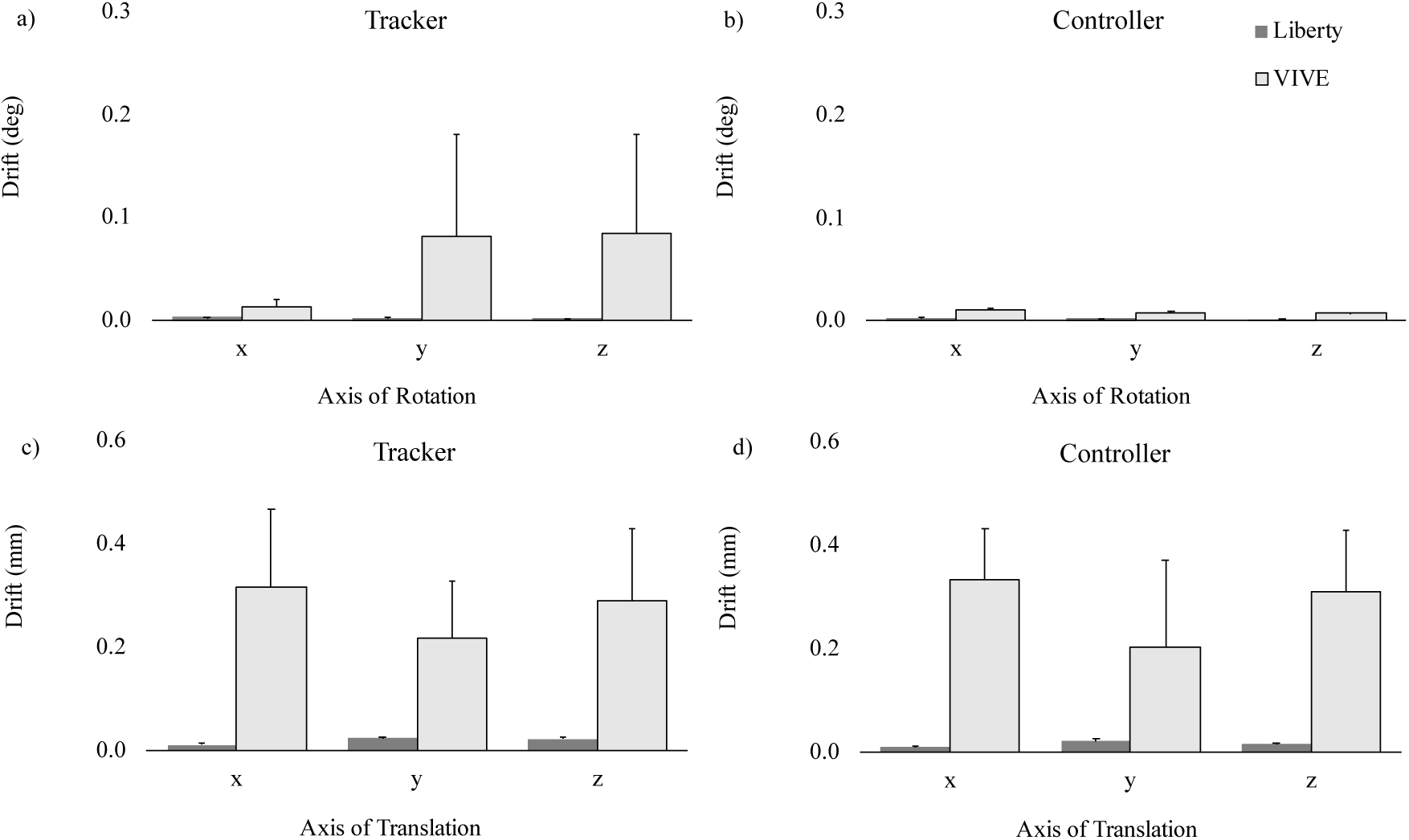
Comparison of rotational (a and b) and translational (c and d) drift measurements (mean ± SD) between the VIVE tracker and the Liberty sensor (a and c) and the VIVE controller and Liberty sensor (b and d).

## 4. Discussion

This study aimed to compare measurements of static rotations and translations between the VIVE handheld controller and tracker to a Liberty sensor, and to quantify rotational and translational drift in all three sensors. These measurements were meant to inform whether the VIVE controller and tracker are accurate enough in measuring position and orientation to use for the collection of kinematic data.

The error measurements of both VIVE sensors were very low, with all mean rotational errors falling below 0.4° and mean translational errors below 3 mm. In all instances when the rotation or translation increment was larger than zero, percent error was less than 0.1%. Rotational error across all three axes appeared to be consistent when using the controller. However, when using the tracker, error about the y-axis appeared to be higher than about the x and z-axes. Even with this inconsistency, mean error about the y-axis of the tracker was only 0.2°. Conversely, translational errors appeared consistent across all axes when using the tracker, and when using the controller error along the z-axis appeared to be lower than along the x and y-axes. Even with these slight variations between axes, all rotational and translational components of the controller and the tracker aligned very closely with the Liberty measurements. These errors fell well within the range of normalcy for research grade systems, which regularly report translational accuracy between 1.0 mm and 2.0 mm and rotational accuracy between 0.5 ° and 1.0 ° but often demonstrate much higher errors in practice (Frantz et al., 2003; Nafis et al., 2006).

Drift measurements throughout the 10 second capture periods from both VIVE sensors were consistently higher in all components than drift from the Liberty sensor. However, all mean rotational drift measures fell below 0.1° and all translational measures below 0.35 mm. The lowest measures of drift by the VIVE system were seen in the controller’s rotational components, while the tracker displayed slightly higher measures of mean rotational drift. As these sensors use embedded IMUs, it is expected that drift would accumulate over time. However, the light-emitting boxes and optical sensors integrate with the output from the IMU to provide 60 drift corrections per second. This mechanism prevents accumulation of drift in the signal.

These slight differences between the VIVE tracker, VIVE controller, and Liberty sensor may be important to consider when making kinematic measurements over very small ranges of motion. Studies involving surgical tasks, for example, which require minimum system accuracy of 1.5 mm, may want to consider using other systems (Birkfellner et al., 1998). However, for larger movements we believe that all of these sensors are accurate enough for the collection of kinematic data. This study was performed using a single VR system, and therefore cannot be guaranteed to represent all systems of the same model. However, this practice is common in validation studies and the protocol is replicable a wider scale (Nafis et al., 2006). Additionally, these data are limited in that they are static measurements taken in the center of the calibrated space; further testing on temporal signal components and the entirety of the space would be necessary for studies with a strong timing requirement or those which plan to utilize all of the calibrated volume. Further, in comparison to a traditional motion capture system, the VIVE sensors are larger and more difficult to affix to segments. The main considerations when choosing between these sensors should therefore be study design, portability, and cost.

